# Exploring DNA methylation profiles in the pathogenesis of human osteoporosis via whole-genome bisulfite sequencing

**DOI:** 10.64898/2026.01.05.697801

**Authors:** Yinyin Zhang, Guoying Wu, Jialu Hou, Yeling Zhong, Yukai Zhang, Shishuo Xiong, Zehua Guo, Ying Li

**Author notes:** Corresponding author: Ying Li. These authors contributed equally: Author Yinyin Zhang.

## Abstract

**Background:** Osteoporosis is a prevalent bone metabolic disorder characterized by reduced bone mass, disruption of bone microarchitecture, and increased bone fragility, leading to a heightened risk of fracture. This condition significantly impairs patients’ quality of life and increases mortality risk. Emerging evidence suggests that DNA methylation may play a crucial role in regulating the expression of genes related to bone metabolism, thereby influencing the development of osteoporosis. However, the precise relationship between DNA methylation and osteoporosis remains unclear and warrants further investigation.

**Results:** Our study revealed significant differences in both the quantity and ratio of DNA methylation between individuals with osteoporosis and non-osteoporosis controls, with differences predominantly occurring in CpG islands. GO/KEGG enrichment analyses highlighted distinct osteoporosis-related gene pathways. Notably, we identified six genes, MSX1, HOXD4, AXIN2, WNT5A, TGFB1, and STAT3, respectively, that are potentially involved in the pathogenesis of osteoporosis and are broadly involved in various diseases and biological processes.

**Conclusions:** These findings indicate distinct methylation patterns between osteoporosis patients and healthy individuals, with differential methylation levels in genes associated with osteoporosis. This research offers new insights into the epigenetic mechanisms underlying osteoporosis.

## Introductions

Osteoporosis is a prevalent metabolic bone disease characterized by diminished bone mass and microarchitectural deterioration, leading to enhanced bone fragility and increased fracture risk. It currently affects an estimated 200 million individuals globally, with approximately one-third of women and one-fifth of men over the age of 50 experiencing osteoporotic fractures. Driven by aging populations and lifestyle factors, its incidence continues to rise, posing a substantial public health burden due to high morbidity, mortality, and associated healthcare costs [1]. Hip fractures represent a particularly severe outcome, associated with a one-year mortality rate of 20–24% and a significant loss of independence; about one-third of survivors become fully dependent or require institutional care within a year post-fracture [2].

The pathogenesis of osteoporosis is multifaceted, involving a complex interplay of genetic, hormonal, and environmental factors. In recent years, epigenetic mechanisms, particularly DNA methylation, have emerged as critical regulators in bone homeostasis and the development of osteoporosis[3]. DNA methylation, the addition of a methyl group to the fifth carbon of cytosine within CpG dinucleotides, is a key heritable epigenetic modification that modulates gene expression without altering the underlying DNA sequence. This process is essential for governing fundamental biological programs such as embryonic development, genomic imprinting, and cellular differentiation [4].

Aberrant DNA methylation patterns are implicated in a wide array of pathologies, including cancer, neurological disorders, and metabolic diseases like osteoporosis [5, 6]. In bone biology, altered methylation can dysregulate the expression of genes pivotal for bone formation and resorption. For instance, promoter hypermethylation of osteogenic genes may suppress their expression, impairing osteoblast function, while hypomethylation of genes promoting osteoclastogenesis can exacerbate bone loss[7–9]. Thus, DNA methylation serves as a vital layer of regulation for bone stability, and its dysregulation appears integral to the pathogenesis of osteoporosis.

Furthermore, DNA methylation profiles hold promise as potential biomarkers for early risk stratification, diagnosis, and prognosis. Epigenome-wide association studies (EWAS) have identified specific differentially methylated regions (DMRs) correlated with bone mineral density (BMD) and fracture risk [10]. Such epigenetic signatures could facilitate the identification of high-risk individuals and inform personalized therapeutic strategies. Additionally, the reversible nature of DNA methylation opens novel therapeutic avenues. Epigenetic drugs, such as DNA methyltransferase inhibitors, which aim to restore normative gene expression patterns, have shown potential in modulating bone metabolism [11].

To comprehensively map the DNA methylome, whole-genome bisulfite sequencing (WGBS) has become a powerful high-throughput technology. WGBS combines bisulfite conversion of DNA with next-generation sequencing to achieve single-base-pair resolution of methylation states across the entire genome [12]. During bisulfite treatment, unmethylated cytosines are deaminated to uracil, while methylated cytosines remain unchanged. This allows for the precise quantification of methylation levels at CpG sites as well as in non-CpG contexts [13].

This study employed WGBS to perform a genome-wide analysis of DNA methylation in individuals with and without osteoporosis. By systematically comparing differential methylation patterns, we aimed to identify genes and regulatory pathways closely associated with osteoporosis, thereby elucidating the role of epigenetic dysregulation in its etiology.

## Methods

### Ethical Statement

This study was conducted in accordance with the Declaration of Helsinki and was approved by the Ethics Committee of The Third Affiliated Hospital of Guangzhou University of Chinese Medicine (Approval No. 20240531-009, Date: 01/06/2024). The ethical approval is valid from 01/06/2024 to 01/06/2025. Participant recruitment took place from 05/06/2024 to 20/11/2024. All participants, or their legal guardians, provided written informed consent prior to enrollment in the study.

### Study Subjects and Sample Collection

This case-control study enrolled 20 participants aged 50–75 years, including 10 patients diagnosed with osteoporosis and 10 non-osteoporotic controls. All participants provided written informed consent prior to enrollment, and the study protocol was approved by the appropriate institutional review board. Participants were categorized into two groups: the osteoporosis group (O group, n=10) and the non-osteoporosis control group (N group, n=10). Bone mineral density (BMD) was measured for all subjects using dual-energy X-ray absorptiometry (DXA; Hologic Discovery, USA). Age and body mass index (BMI) were matched between groups to ensure comparability. Bone tissue specimens were obtained during elective lumbar spinal surgery. After dissection to remove adjacent soft tissues, samples were immediately snap-frozen in liquid nitrogen and stored at –80°C until DNA extraction **(Figure 1)**.

**Figure 1.**
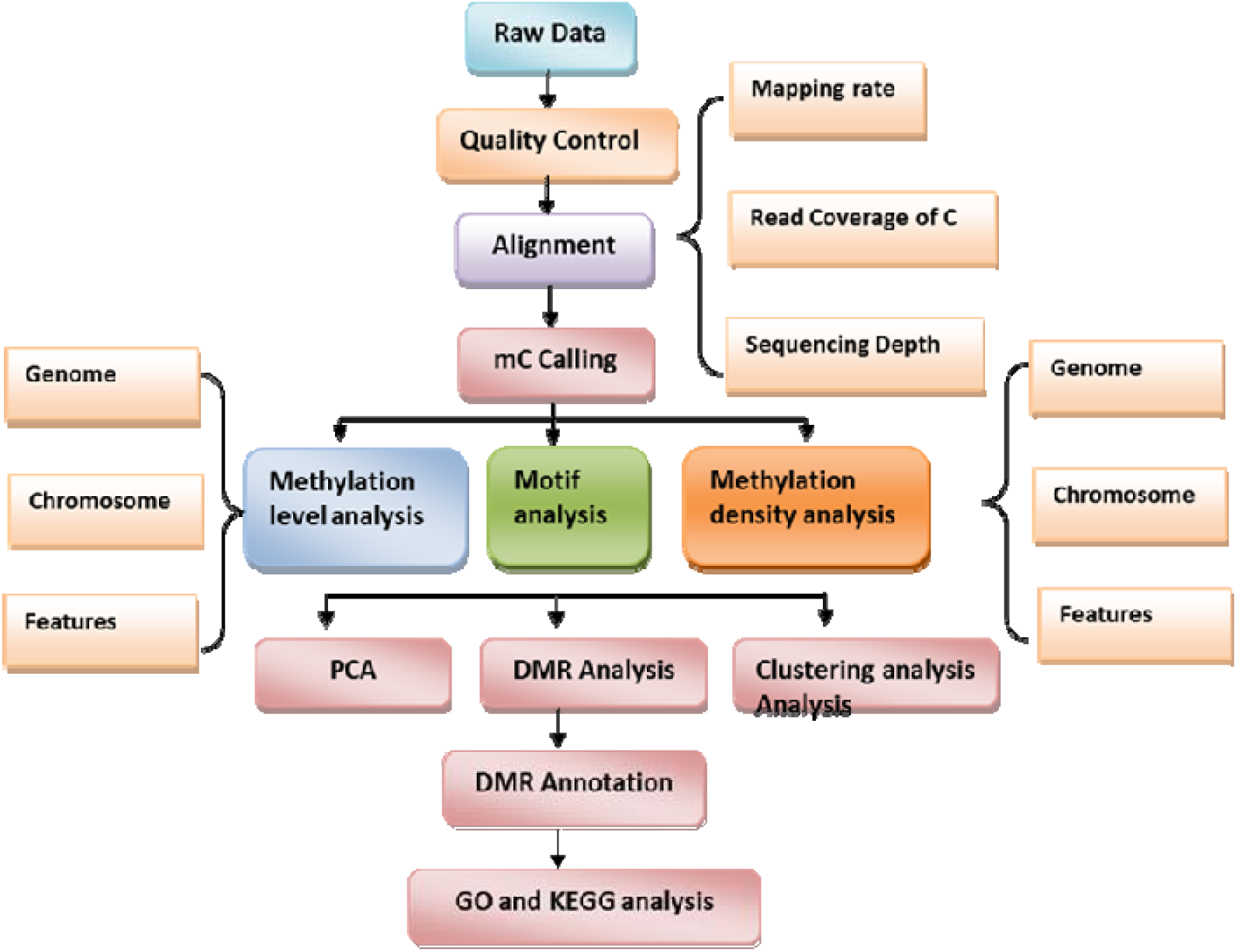
Whole-genome bisulfite sequencing analysis flowchart.

### DNA Extraction from Bone Tissue

Bone samples were rinsed with deionized water and a mild detergent to remove surface contaminants. A sterile surgical blade was used to remove a thin outer layer, exposing the inner bone matrix. Tissues were flash-frozen in liquid nitrogen and mechanically pulverized into a fine powder. Bone powder was decalcified in 0.5 M EDTA (pH 8.0) at 4°C for 72 hours with gentle agitation. Decalcified tissue was subsequently digested in lysis buffer, and genomic DNA was isolated using a commercial DNA extraction kit according to the manufacturer’s instructions. DNA purity and concentration were assessed using a NanoDrop spectrophotometer and Qubit fluorometer, respectively.

### Library Preparation and Whole-Genome Bisulfite Sequencing

Genomic DNA was fragmented by sonication, followed by end repair, 3′ adenylation, and ligation of methylated adapters. Adapter-ligated DNA underwent bisulfite conversion using the EZ DNA Methylation-Gold Kit. Converted libraries were amplified by PCR, purified, and assessed for quality using an Agilent 2100 Bioanalyzer. Quantification was performed with a Qubit® 2.0 Fluorometer. Pooled libraries were loaded onto an Illumina NovaSeq 6000 platform for 150 bp paired-end sequencing, following standard Illumina protocols. Sequencing metrics, including cluster density and phasing rates, were monitored in real time **(Table 1)**.

**Table 1.** Reagents for library construction and quality inspection.

| Product/Reagent | Manufacturer | Catalog No. |
| --- | --- | --- |
| SureSelectXT Methyl Reagent kit, HSQ,16 | Agilent | G9651A |
| EZ-DNA Methylation-Gold Kit | Zymo | D5006 |
| SureSelectXT Human Methyl-Seq Capture Library | Agilent | 5190-4661 |
| ExKubit dsDNA HS Analysis Kit | Excell | NGS00-3012 |
| Agilent High Sensitivity DNA Kit | Agilent | 5067-4626 |

### Sequencing Data Quality Control

Raw sequencing reads were evaluated for quality using FastQC. Base calling quality was expressed as Q-scores, where Q = –10 log□□(E) and E represents the estimated error probability. Due to bisulfite conversion of unmethylated cytosines to thymines, a reduction in C-content and a corresponding increase in T-content were expected and confirmed. Per-sample sequencing output targeted ∼20 Gb, with >90% of bases achieving Q ≥ 20 (error rate < 1%).

### Read Preprocessing and Alignment

Adapter sequences and low-quality bases were trimmed using Trim Galore (v0.4.1) with the following criteria: removal of reads with >50% of bases having Phred score < 20, trimming of 3′-end bases with Q < 20, and discarding reads shorter than 70 bp after trimming. Cleaned reads were aligned to the human reference genome (GRCh37/hg19) using Bismark (v0.15.0) in conjunction with Bowtie 2 (v2.2.9)[14]. PCR duplicates were removed using the deduplicate_bismark tool. Mapping efficiency, coverage depth, and bisulfite conversion rates were calculated for each sample.

### Methylation Calling and Differential Analysis

Cytosine methylation levels were extracted using bismark_methylation extractor[15]. Methylation ratios (mC/C) were calculated for each CpG site covered by at least 5 reads. Genome-wide methylation profiles were summarized and visualized using the R package methylKit (v0.9.5)[16, 17]. Differentially methylated regions (DMRs) were identified with the DSS package using a smoothing-based approach. Regions with |Δβ| > 0.1 (10% methylation difference) and an adjusted p-value < 0.05 (FDR correction) were considered significant. CpG islands were annotated using the annotatr package in R based on UCSC definitions.

### Functional Enrichment Analysis

Genes associated with significant DMRs were subjected to functional annotation. Gene Ontology (GO) and Kyoto Encyclopedia of Genes and Genomes (KEGG) pathway enrichment analyses were performed using the clusterProfiler package in R. A hypergeometric test was applied, with Bonferroni-corrected p-values ≤ 0.05 considered statistically significant.

### Protein-Protein Interaction Network Analysis

To investigate the functional associations among the genes associated with differentially methylated regions (DMRs), a protein-protein interaction (PPI) network was constructed using the STRING database (version 12.0; https://string-db.org). The list of DMR-associated genes was submitted to STRING, and interactions were retrieved with a minimum required interaction score set to 0.400 (medium confidence). The resulting network was visualized within the STRING platform, and hub genes were identified based on their connectivity degree. This analysis aimed to reveal potential core regulatory modules and functional clusters among the epigenetically dysregulated genes in osteoporosis.

### Statistical Analysis

Clinical and demographic variables were analyzed using R 4.5.2. Continuous variables conforming to normal distribution are presented as mean ± standard deviation and compared via independent two-sample t-tests. Categorical variables are expressed as counts and compared using chi-square tests. Non-normally distributed data were analyzed with nonparametric tests. Spearman’s rank correlation was used to assess associations between methylation levels and clinical parameters. A two-tailed p-value < 0.05 was considered statistically significant.

## Results

### Baseline Characteristics of the Study Cohort

A total of 20 participants were enrolled in this case-control study and categorized into two groups: the osteoporosis group (O group, n=10) and the non-osteoporosis control group (N group, n=10).

The two groups were well-matched at baseline. The mean age was 65.20 (±4.44) years in the O group and 63.60 (±9.37) years in the N group (P=0.85). There were no statistically significant differences in height (157.80 ± 8.66 cm vs. 161.20 ± 10.81 cm, P=0.448), weight (60.10 ± 8.48 kg vs. 62.30 ± 10.89 kg, P=0.62), or body mass index (23.69 ± 3.22 kg/m² vs. 23.94 ± 3.19 kg/m², P=0.86) between the O and N groups. The cohort was predominantly female (85%), with a comparable sex distribution between groups (80% female in O group vs. 90% in N group, P>0.99)**(Table 2)**. These results confirm that the two groups were comparable in terms of major baseline characteristics, minimizing potential confounding effects in the subsequent epigenetic analysis.

**Table 2.** Demographic and clinical characteristics at baseline.

| Variable | Overall, n = 20<sup>1</sup> | N group n = 10 (25%)<sup>1</sup> | O group N = 10 (75%)<sup>1</sup> | P-value<sup>2</sup> |
| --- | --- | --- | --- | --- |
| Age(years) | 64.40 ± 7.18 | 63.60 ± (9.37) | 65.20 ± (4.44) | 0.85 |
| Height(cm) | 159.50± (9.69) | 161.20 ± (10.81) | 157.80 ± (8.66) | 0.448 |
| Weight(kg) | 61.20± (9.57) | 62.30 ± (10.89) | 60.10 ± (8.48) | 0.62 |
| BMI(kg/m<sup>2</sup>) | 23.81± (3.12) | 23.94± (3.19) | 23.69 ± (3.22) | 0.86 |
| **Sex** |  |  |  |  |
| Female ,No. (%) | 17 (85.00%) | 9 (90.00%) | 8 (80.00%) | >0.99 |
| Male.No. (%) | 3 (15.00%) | 1 (10.00%) | 2 (20.00%) |  |

<sup>1</sup>n (%); Mean ± SD; Median [IQR], <sup>2</sup>Fisher's exact test; Two Sample t-test; Wilcoxon rank sum test

### Genome-wide mapping of DNA methylation and genome coverage statistics

To comprehensively profile genome-wide DNA methylation differences in bone tissue between osteoporotic patients and healthy controls, we conducted whole-genome bisulfite sequencing on a cohort of 20 clinical specimens. Following stringent quality control and filtering, high-quality sequencing data were obtained for all samples in both the osteoporosis group (O group, n=10) and the normal control group (N group, n=10). Each sample yielded approximately 20 Gb of clean data with 150-base pair paired-end reads. Subsequent alignment and quality assessment demonstrated that all samples achieved a mapping efficiency exceeding 80% and a genome coverage breadth of over 99%. The average sequencing depth across the genome was consistently maintained at ≥150-fold for all samples, ensuring robust detection of methylation states **(Table 3)**

**Table 3.** Whole genome methylation sequencing data.

| Group | Clean base | Clean reads | Mapped (%) | Base coverage rate (%) | Mean depth |
| --- | --- | --- | --- | --- | --- |
| O | 28.35±7.55 | 189126106±50348380 | 84.81±2.01 | 99.48±0.02 | 201.55±60.72 |
| N | 21.04±1.90 | 139598858±13666689 | 81.71±0.52 | 99.53±0.02 | 153.36±15.80 |

O: osteoporosis; N: non-osteoporosis; Clean base: Clean data after sequencing; Clean reads: number of reads that have been filtered; Mapped (%): ratio of reads to the Clean reads on the only match; Base coverage rate (%): the percentage of bases in a genome or target region that are covered by at least one sequencing read, indicating the
extent of the genome that has been successfully sequenced; Mean depth: the average sequencing depth of each site in the captured region.

**Table 4.** Average methylation levels in different backgrounds.

| Group | MeanCG(%) | MeanCHG(%) | MeanCHH(%) |
| --- | --- | --- | --- |
| O | 48.99 ± 2.46* | 2.41 ± 0.08 | 2.23 ± 0.06 |
| N | 46.84 ± 1.01 | 2.17 ± 0.04 | 2.04 ± 0.06 |

Data represent means ± SD (n = 10 samples replicates for O and N group), \*refers as significant difference ( $P < 0.05$ ). MeanCG(%): average methylation levels in CG regions, mCG counts/ (CG counts on the genome)\*100%; MeanCHG(%): average methylation levels in CHG region, mCHG counts/ (CHG counts on the genome)\*100%; MeanCHH(%): average methylation levels in CHH region, mCHH counts/ (CHH counts on the genome)

### Methylation level and distribution characteristics

To compare the genome-wide distribution of DNA methylation contexts between osteoporotic and healthy bone tissues, we analyzed the proportion of methylated cytosines in CG, CHG, and CHH sequence contexts. In the osteoporosis group, methylation occurred predominantly in the CG context (91.35%), with minor fractions in CHG (4.49%) and CHH (4.16%). Similarly, the normal control group showed 91.75% CG, 4.25% CHG, and 4.00% CHH methylation **(Figure 2A)**, confirming CG methylation as the dominant form in both groups.

**Figure 2.**
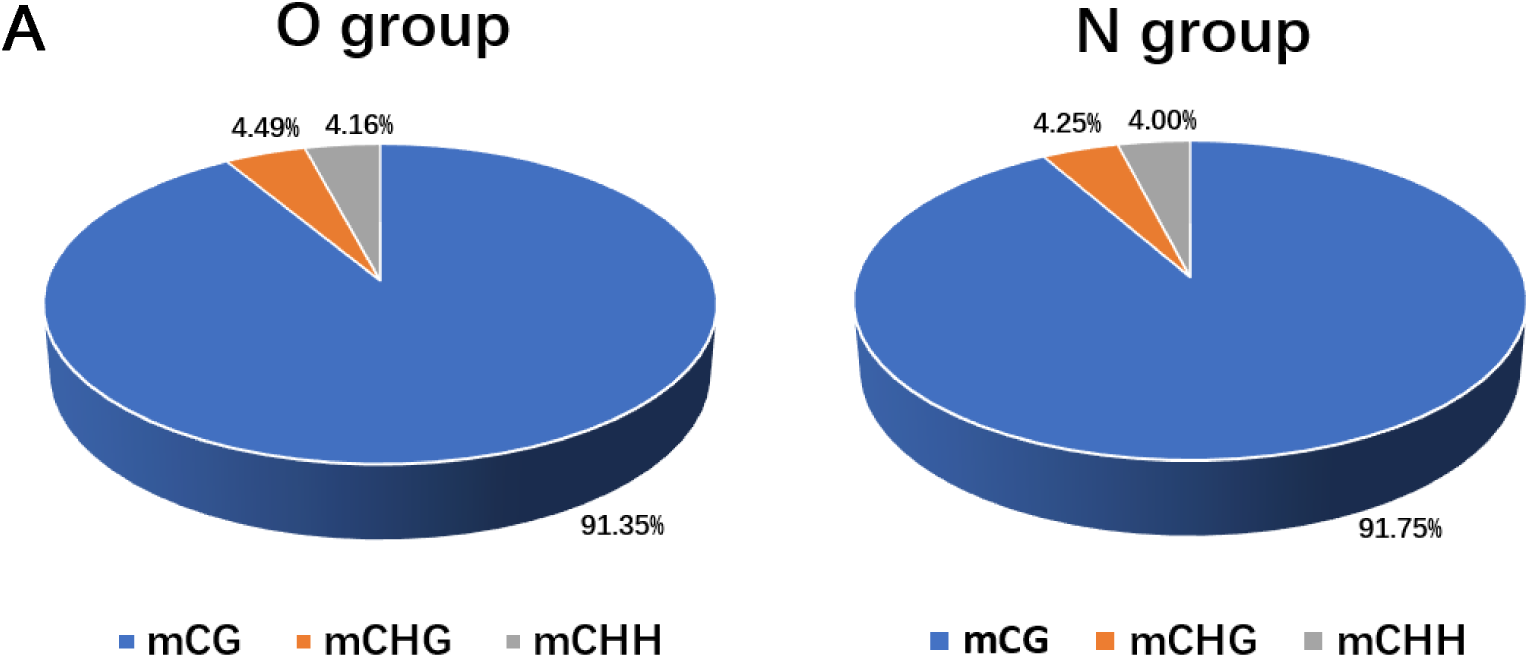
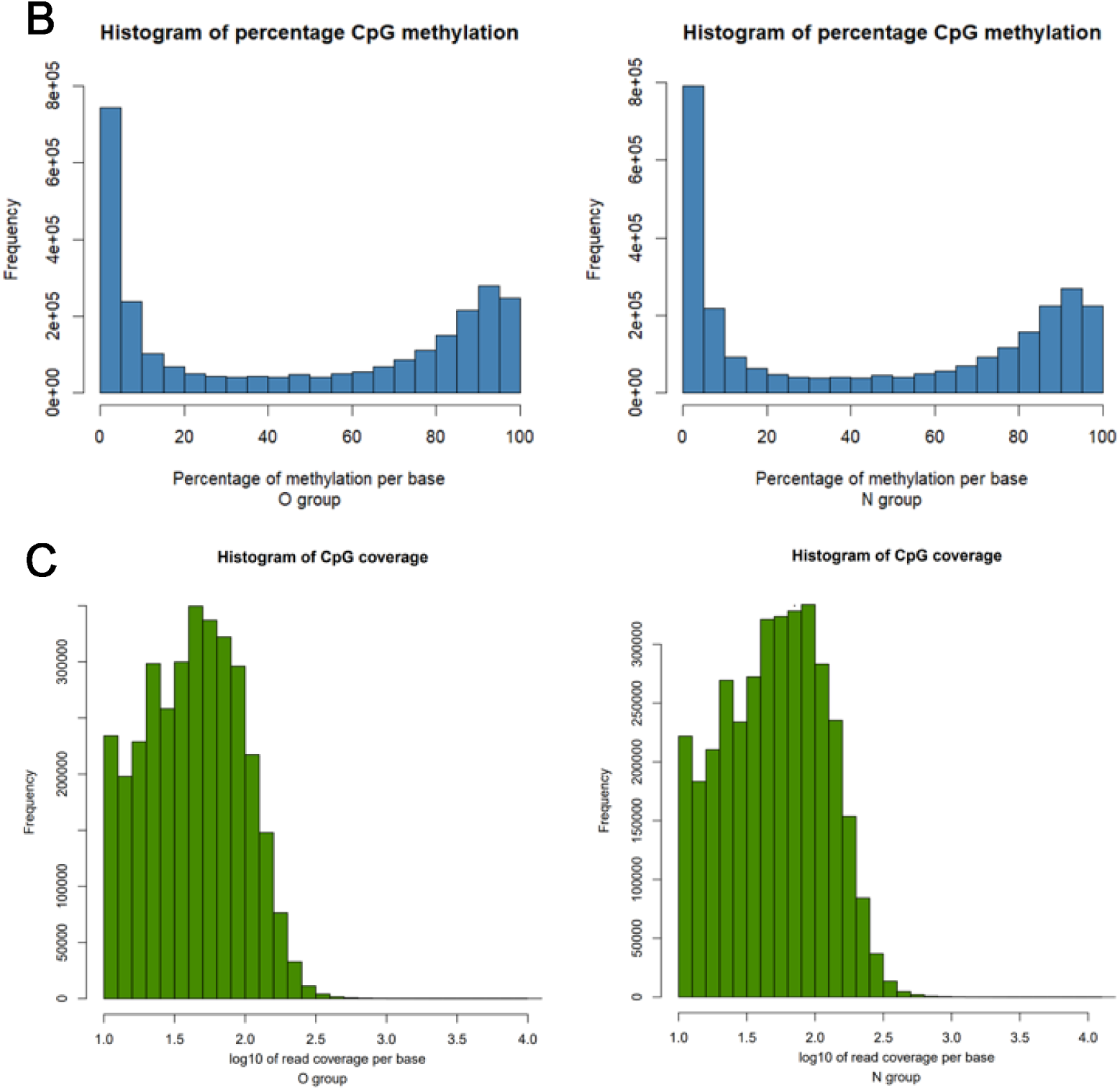
The average ratio and distribution characteristics of group O and Group N DNA methylation types. **(A)** Methylation ratio in different backgrounds. O group, osteoporosis group. N group, normal group. The blue, orange and gray colors represent mCG, mCHG and mCHH, respectively. **(B)** Average percentage of single base methylation probability per group. The horizontal coordinate indicates the probability of methylation, and the vertical coordinate indicates the number of CpG sites for this degree of methylation. The probability of methylation is determined by the amount of C measured at each site/the amount of (C+T) measured at that. **(C)** The coverage depth of methylation sites was detected in each CpG background. The abscissa represents the coverage of each base, and the ordinate represents the proportion of CpG under that coverage.

We next quantified methylation levels in each sequence context using stringent bisulfite conversion efficiency thresholds. The osteoporosis group exhibited methylation levels of 48.99 ± 2.46% (CG), 2.41 ± 0.08% (CHG), and 2.23 ± 0.06% (CHH), while the normal group showed 46.84 ± 1.01% (CG), 2.17 ± 0.04% (CHG), and 2.04 ± 0.06% (CHH) **(Table 3)**. A statistically significant difference between groups was observed specifically in CG methylation (P < 0.05).

To further characterize CG methylation patterns, we generated dual-parameter frequency histograms assessing both methylation probability and coverage depth. Methylation probability distribution revealed a bimodal pattern in both groups, with peaks at <10% and >90% methylation frequencies and few sites at intermediate levels **(Figure 2B)**. Coverage analysis indicated a progressive decrease in the proportion of CpG sites with increasing sequencing depth **(Figure 2C)**. Comparative assessment demonstrated that the osteoporosis group consistently displayed higher methylation probabilities and greater per-site coverage depths than the normal controls across individually mapped cytosine positions.

### Differential DNA methylation regions distribution and characteristics

Volcano plot analysis of DMRs revealed 996 regions with significant methylation alterations (795 hypermethylated and 201 hypomethylated) out of 2,484 analyzed regions **(Figure 3A)**, corresponding to a hyper- to hypomethylation ratio of 3.96:1, demonstrating a genome-wide shift toward hypermethylation in osteoporotic bone.

**Figure 3.**
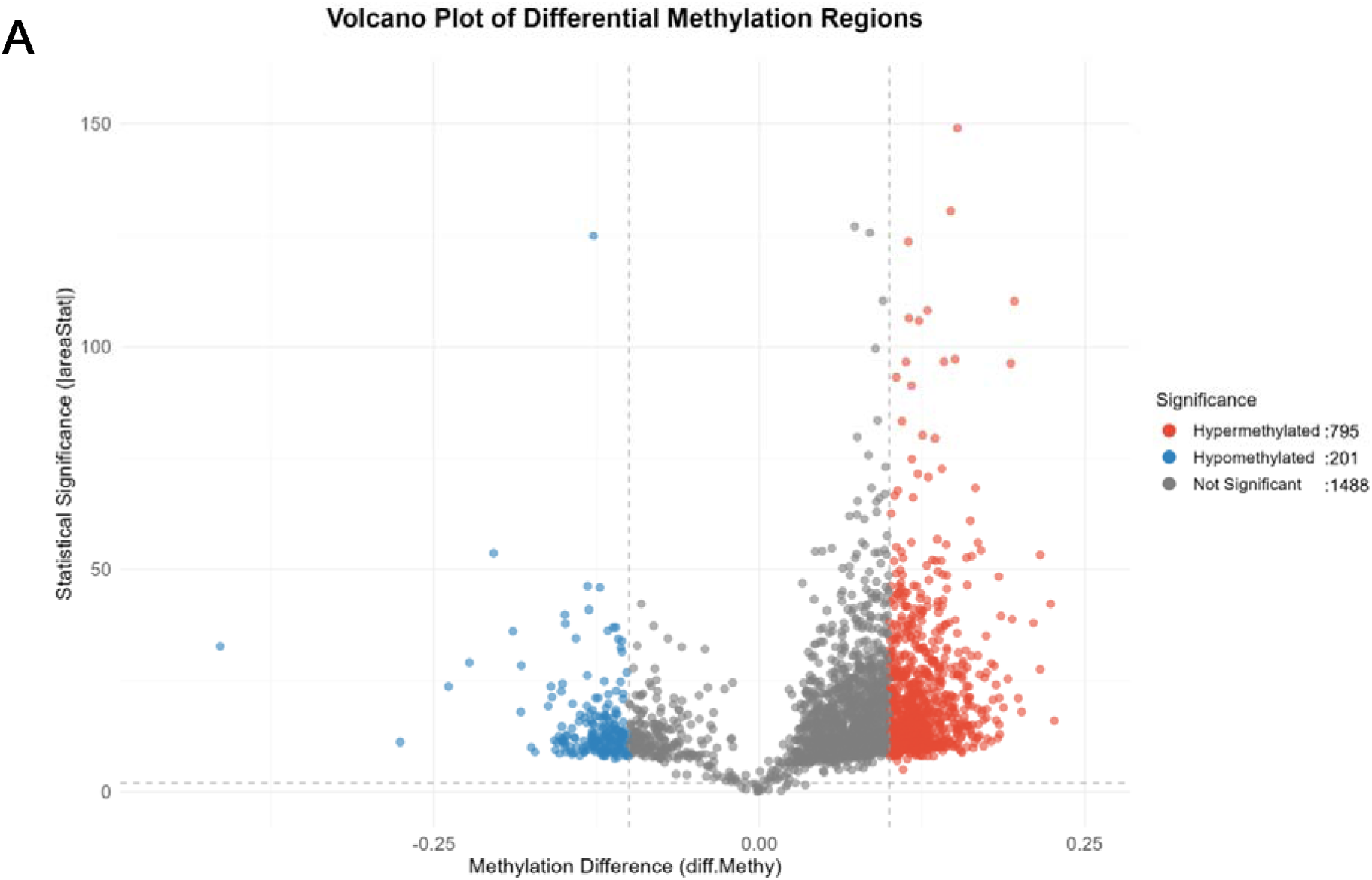
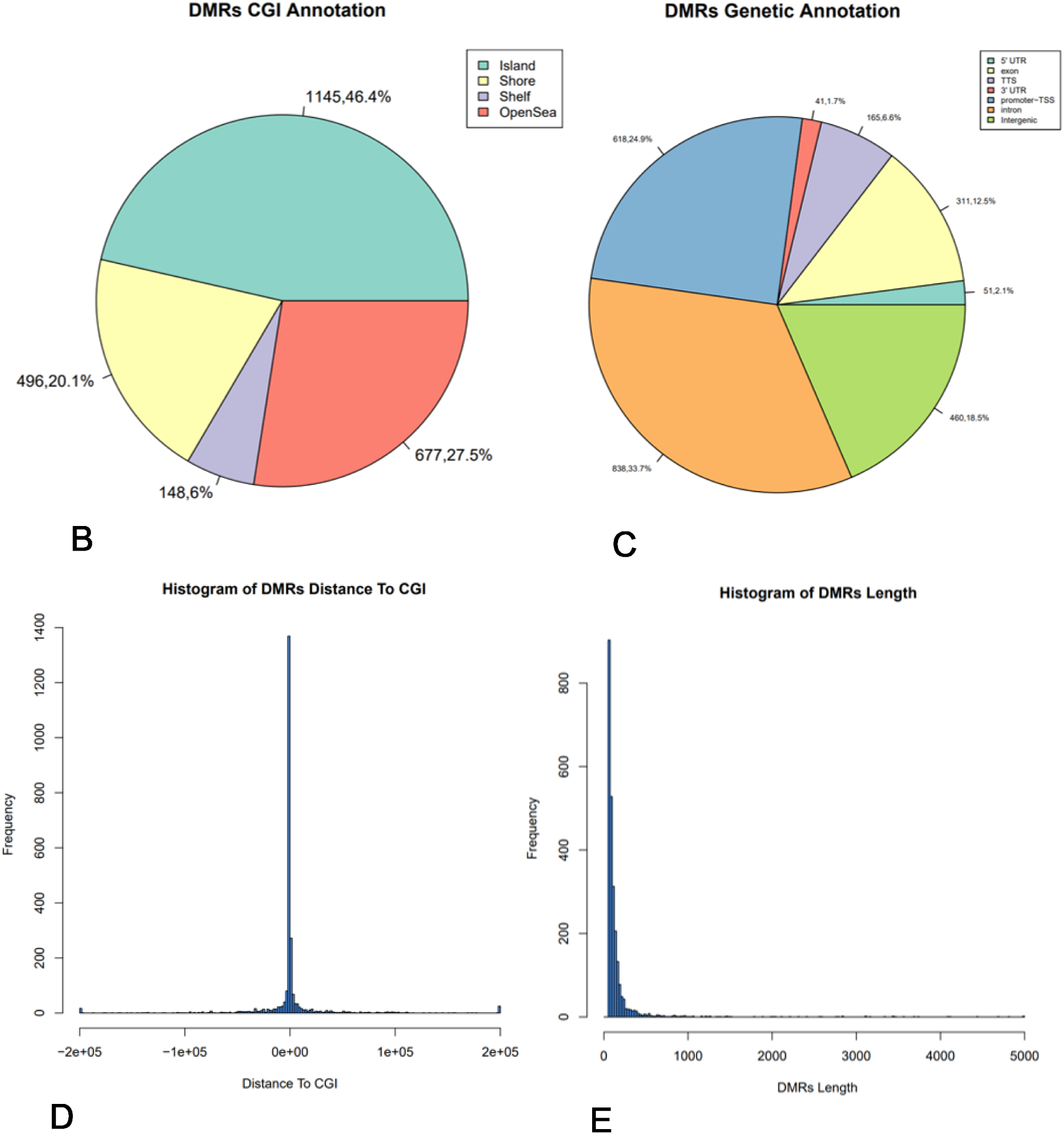
Differential DNA methylation regions distribution and characteristics. **(A)** Distribution of differential methylation regions in CpG islands and different components of CPG islands. Island indicates that DMRs overlaps CGI; Shore indicates that the distance between DMRs and CGI is between 1-2kb. Shelf indicates a distance between 2 and 4kb. OpenSea indicates that the distance between the two is greater than 4kb. **(B)** Distribution of differential methylation regions on different gene elements. **(C)** The frequency of the difference between methyl region segments and CGI relative positions. The horizontal coordinate refers to the upstream and downstream position of DMRs relative to CGI, and the vertical coordinate refers to the frequency of differential methylation segments at specific locations. **(D)** Length distribution of differential methylation regions. The horizontal coordinate refers to the length of DMRs, and the total left side refers to the frequency of DMRs for that length.

Genomic annotation of DMRs indicated their predominant enrichment within CpG islands (CGIs) (1,145 regions, 46.4%), with decreasing density in CGI shores (496 regions, 20.1%) and shelves (407 regions, 16.4%) **(Figure 3B)**. Functional compartment analysis further showed that DMRs were significantly enriched in intronic regions (33.7%) and promoter/enhancer elements (24.9%), while being minimally represented in 5′ UTR (2.1%) and 3′ UTR regions (1.7%) **(Figure 3C)**. Distance-frequency distribution analysis relative to CGIs confirmed an inverse relationship between DMR abundance and genomic distance from CGIs **(Figure 3D)**. To characterize DMR architecture, we analyzed the distribution of region lengths. Length-frequency histograms revealed a strong negative correlation between DMR length and abundance, with over 90% of DMRs spanning less than 1,000 bp **(Figure 3E)**.

### GO and KEGG enrichment

To elucidate the biological functions of the DMRs identified between osteoporotic patients and healthy individuals, pathway enrichment analyses were conducted based on the GO and KEGG databases. In addition, PPI network analysis was performed using the STRING database. A total of 996 significant DMRs were analyzed, and the top 10 regions showing the most pronounced differences, along with osteoporosis-related GO terms and KEGG pathways, are presented.

In the GO enrichment analysis, DMRs were significantly enriched in biological processes such as gland development, epithelial cell proliferation, embryonic organ development, and embryonic organ morphogenesis **(Figure 4A)**. KEGG pathway analysis revealed significant enrichment in pathways including non-small cell lung cancer, chronic myeloid leukemia, and parathyroid hormone synthesis **(Figure 4B)**.

**Figure 4.**
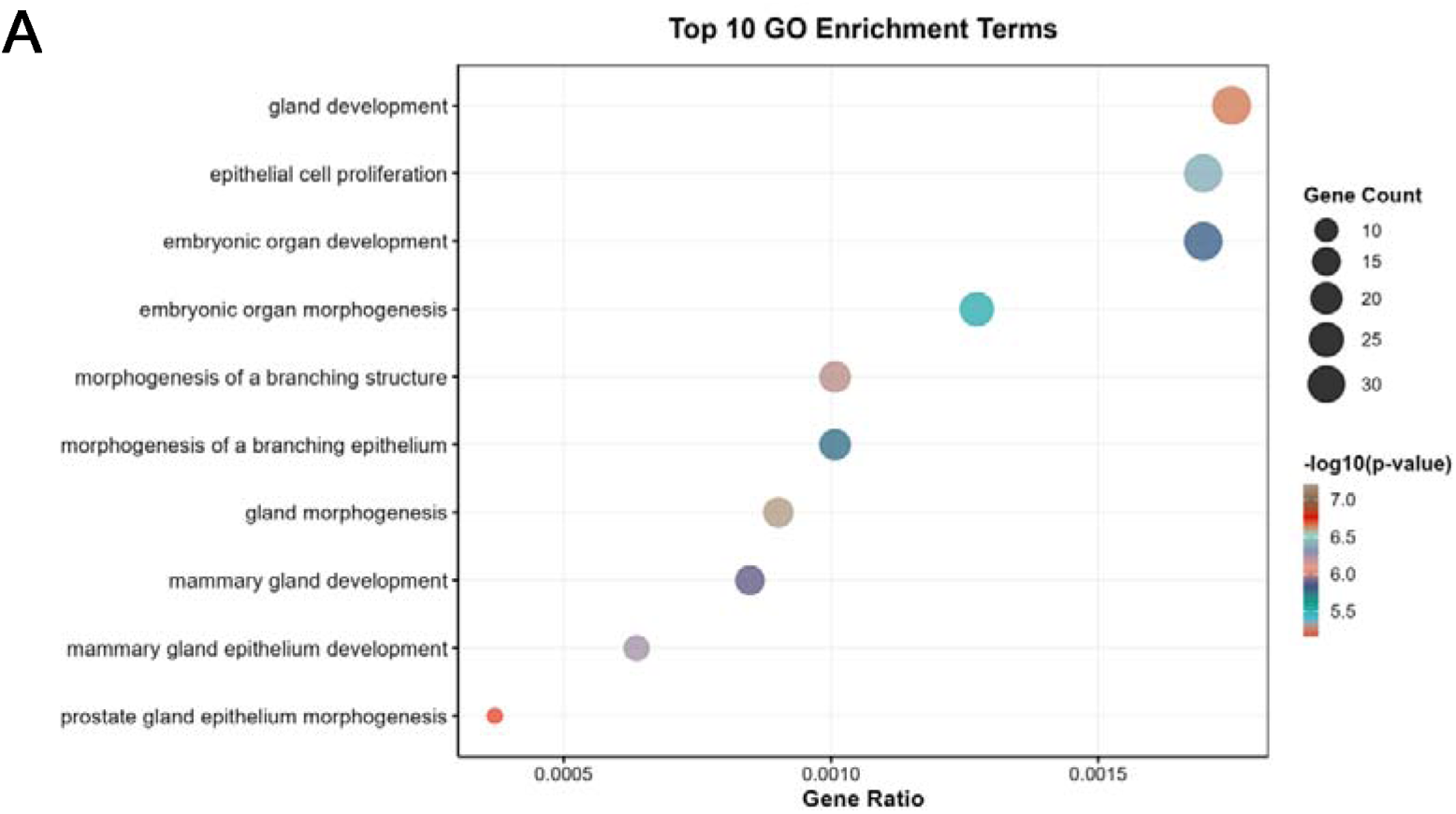
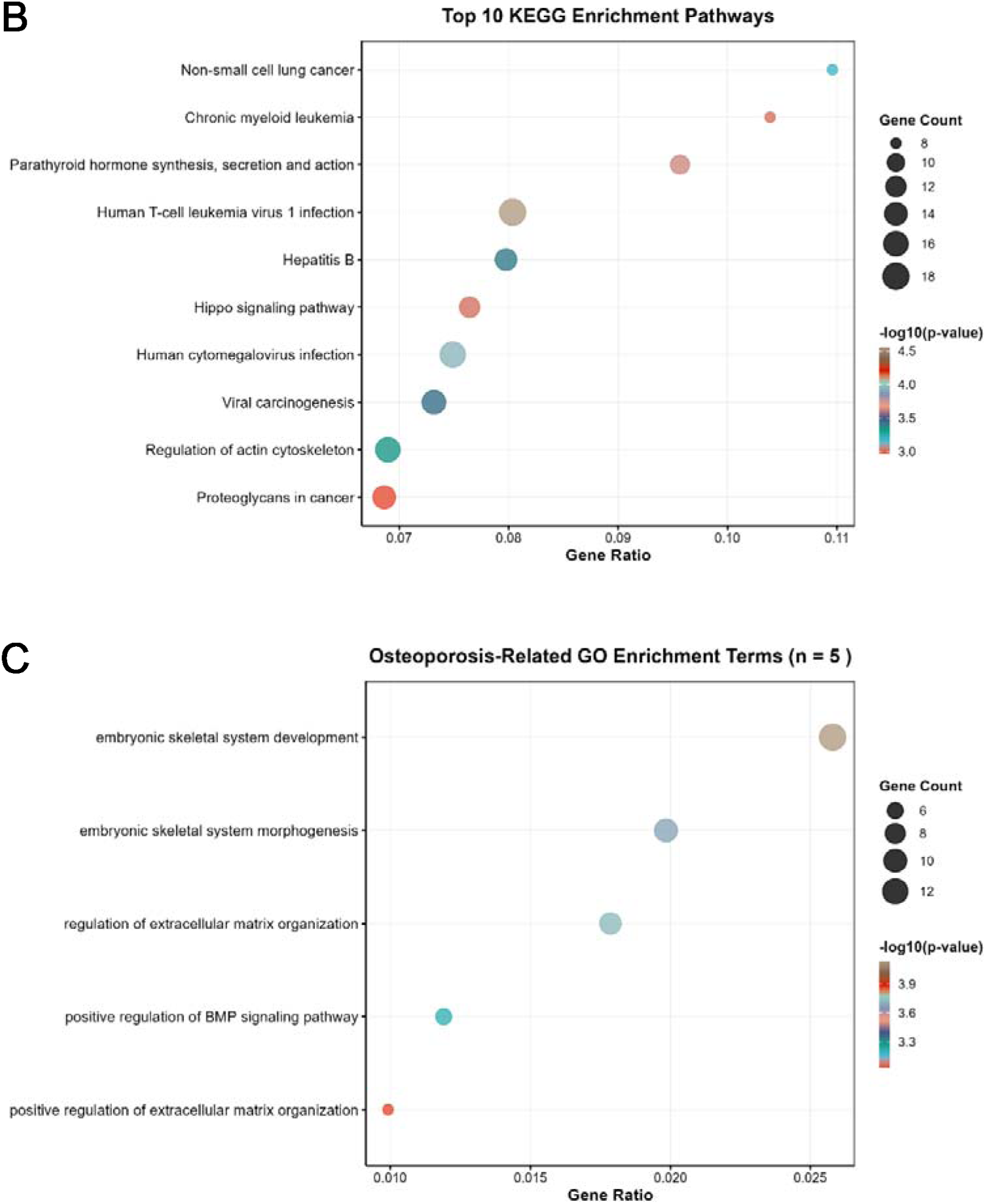
Functional enrichment analysis of DMRs between osteoporotic patients and healthy controls. **(A)** GO enrichment analysis of biological processes. The bubble chart displays significantly enriched GO terms, with the bubble size representing the number of genes or DMRs mapped, and the color intensity indicating the statistical significance. Representative terms include gland development and epithelial cell proliferation. **(B)** KEGG pathway enrichment analysis. The bubble chart illustrates significantly enriched KEGG pathways, where the size and color of the bubbles denote the gene count and the level of enrichment significance, respectively. Key pathways identified include non-small cell lung cancer and chronic myeloid leukemia. **(C)** Screening of osteoporosis-related GO terms via keyword filtering. This bubble chart presents the five GO terms identified as relevant to bone metabolism and extracellular matrix organization, with the bubble size and color similarly reflecting gene counts and statistical significance.

GO entries containing keywords associated with bone biology—such as “bone,” “osteoclast,” “osteoblast,” “ossification,” “mineral,” “cartilage,” “chondrocyte,” “skeletal,” “calcium,” “Wnt,” “BMP,” “TGF,” “collagen,” “extracellular matrix,” and “ECM”—were screened. This search identified five GO terms related to osteoporosis: “embryonic skeletal system development”,“embryonic skeletal system morphogenesis,” “regulation of extracellular matrix organization,” “positive regulation of BMP signaling pathway,” and “positive regulation of extracellular matrix organization” **(Figure 4C)**. Notably, no KEGG pathways directly associated with osteoporosis were identified.

### Differential gene screening and validation in public databases

Through analysis of osteoporosis-related pathways, we identified 19 genes and constructed a PPI network for these genes. Genes with node degrees ≥3 were preliminarily screened, resulting in 11 candidate genes: MSX1, HOXD4, AXIN2, WNT5A, TGFB1, STAT3, RUNX2, NOTCH1, HOXD9, MSX2, BMP7, and HOXD3**(Figure 5)**.

**Figure 5.**
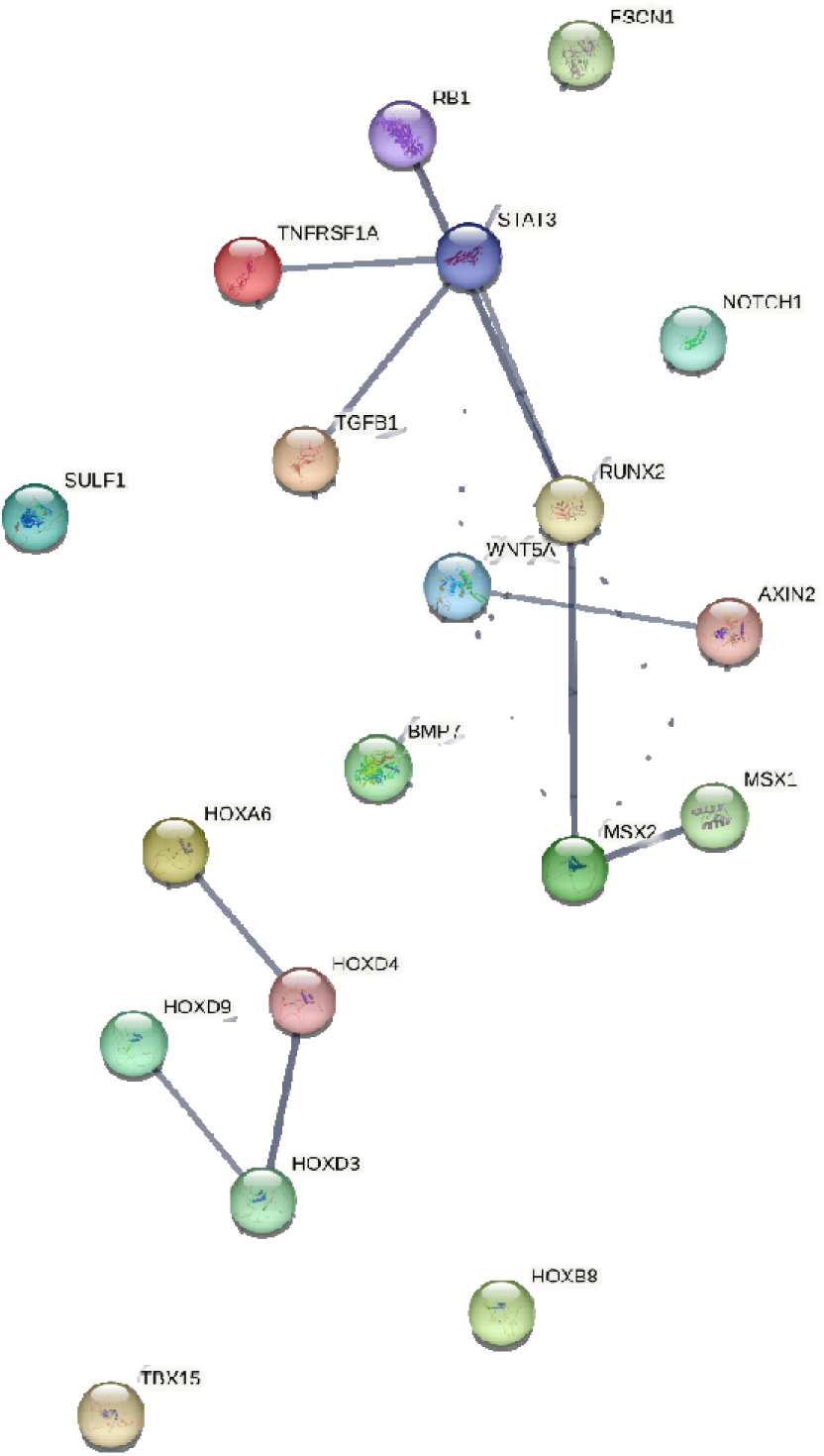
Construction of differential methylation gene networks related to bone development. Analysis of the interaction between DMRs related to muscle development using STRING software according to the interplay index (confidence > 0.4). The interplay index between genes was represented by edge width and transparency. Dark and wide edges indicated high confidence.

Subsequently, we selected the dataset GSE230664 from the GEO database, which is also related to osteoporosis research, to validate the expression of these 11 genes. The results showed that six genes—MSX1, HOXD4, AXIN2, WNT5A, TGFB1, and STAT3—exhibited expression trends consistent with the methylation data. However, none of the genes showed statistically significant differences in expression between the two groups**(Table 5)**.

**Table 5.** Validation of differentially expressed genes in the GEO database.

| Gene | Methylation Change ($\Delta\beta$) | Expression Change (logFC) | Expression adj.P.Val | Methylation-Expression Concordance |
| --- | --- | --- | --- | --- |
| MSX1 | +0.155 | -0.750 | 0.348 | Concordant<br>(Hypermethylation/Downregulation) |
| HOXD4 | +0.095 | -0.804 | 0.080 | Concordant<br>(Hypermethylation/Downregulation) |
| AXIN2 | +0.112 | -0.282 | 0.663 | Concordant<br>(Hypermethylation/Downregulation) |
| WNT5A | +0.099 | -0.121 | 0.973 | Concordant<br>(Hypermethylation/Downregulation) |
| TGFB1 | -0.129 | +1.109 | 0.330 | Concordant<br>(Hypomethylation/Upregulation) |
| STAT3 | -0.112 | +1.316 | 0.436 | Concordant<br>(Hypomethylation/Upregulation) |
| RUNX2 | +0.114 | +0.860 | 0.402 | Discordant |
| NOTCH1 | +0.116 | +0.607 | 0.424 | Discordant |
| HOXD9 | +0.129 | +0.581 | 0.653 | Discordant |
| MSX2 | +0.101 | +0.867 | 0.227 | Discordant |
| BMP7 | +0.112 | +0.048 | 0.963 | Discordant |
| HOXD3 | +0.101 | +0.299 | 0.666 | Discordant |

$\Delta\beta$ : Average methylation difference (O group-N group), positive value indicates hypermethylation. logFC: log2 fold change (O group/N group), negative value indicates downregulation.

## Discussion

The concept of epigenetics, introduced by the British biologist Conrad Hal Waddington in 1942[18], describes how interactions between genes and their products influence cellular development without altering the DNA sequence itself. Epigenetic modifications affect gene activity by altering DNA and chromatin chemical modifications, primarily through DNA methylation, histone modification, noncoding RNA regulation, and chromatin remodeling. As research has advanced, epigenetics has been implicated in the onset and progression of various diseases, including cancer, neurological disorders, and metabolic diseases.

DNA methylation is a key epigenetic modification in which a methyl group is added to the 5th carbon atom of cytosine within DNA, forming 5-methylcytosine. This process is catalyzed by DNA methyltransferases (DNMT), primarily at CpG dinucleotides[19]. DNMT1 maintains methylation patterns during DNA replication, transferring methylation marks from the parent strand to the daughter strand. DNMT3A and DNMT3B establish new methylation patterns during development and differentiation, whereas DNMT3L facilitates DNMT3A and DNMT3B but lacks catalytic activity itself[20]. DNA methylation is generally associated with gene silencing, particularly when promoter regions are highly methylated[21], which suppresses transcription. During development, specific methylation patterns change in response to gene expression needs, and methylation helps suppress transposons and repetitive sequences, maintaining genomic stability.

Osteoporosis development is characterized by an imbalance in bone remodeling, with decreased bone formation and increased bone resorption. Factors such as genetics, the environment, hormonal changes, nutrition, and physical activity influence this balance[22]. DNA methylation also plays a critical role in osteoporosis by regulating the expression of genes related to bone metabolism. The methylation status of specific genes can alter osteoblast and osteoclast function, impacting osteoporosis development. For instance[23], hypermethylation of the RUNX2 gene, which is associated with osteoblast differentiation, can inhibit osteoblast function and reduce bone formation. Conversely, hypomethylation of the RANKL gene, which is linked to osteoclast differentiation, can increase osteoclast activity and increase bone resorption.

Our results suggest that DNA methylation is involved in various biological processes, cellular functions, and disease progression. Although this study cannot directly establish a causal relationship between DNA methylation and osteoporosis, evidence points to a significant association between the two. We observed differences in the methylation levels and expression of methylation-related genes between the osteoporosis and non-osteoporosis groups. Enrichment analysis further revealed differential methylation of several genes associated with osteoporosis, indicating that altered methylation levels of certain genes may be associated with this disease.

Through the integrative analysis of whole-genome methylation profiles from bone tissue and an independent transcriptomic dataset (GSE230664), this study identified several potential epigenetically regulated targets, such as MSX1, AXIN2, and TGFB1, which exhibited consistent directions of change in both methylation and expression. However, we also noted that some genes displaying significant differential methylation in our cohort did not show statistically significant expression differences in the public dataset. This incomplete correspondence between epigenetic alterations and transcriptional output is a common phenomenon in complex biological systems and may be attributed to the following interrelated factors.

First, the function of DNA methylation, particularly at regions outside promoter CpG islands—such as enhancers, gene bodies, or silencers—is highly dependent on cell type, differentiation stage, and specific microenvironmental signals[24]. The bone tissue samples used in this study contain heterogeneous cell populations, including osteoblast progenitors, mature osteoblasts, and osteocytes. A key methylation alteration in a specific progenitor subpopulation may have its corresponding expression signal diluted in bulk RNA analysis. For instance, RUNX2, the master regulator of osteogenic differentiation, exhibits stringent stage-specific expression. The methylation changes observed in its promoter region may significantly affect its transcriptional activity only during specific differentiation windows or within particular cell subsets, which is difficult to capture in bulk sequencing data.

Second, gene expression is co-regulated by a multi-layered network, including transcription factors, histone modifications, and non-coding RNAs. DNA methylation represents one layer within this network, and its effects may be compensated for or buffered by other, stronger regulatory layers. Methylation in a gene’s promoter region could be counteracted by a potent enhancer or specific activating histone modifications, thereby maintaining expression stability. Conversely, even if the methylation status remains unchanged, alterations in other regulatory layers may drive expression changes[25]. This network robustness explains why a single methylation change is sometimes insufficient to produce a detectable expression phenotype.

MSX1 and HOXD4 both encode homeobox transcription factors that are crucial for skeletal patterning and osteoblast commitment[26]. Their hypermethylation and reduced expression suggest an early impairment in osteogenic lineage determination. This epigenetic silencing may lead to a diminished osteoprogenitor cell pool or reduced differentiation capacity, a hallmark of age-related bone loss.

Concurrently, the epigenetic repression of AXIN2 and WNT5A points to a coordinated disruption of Wnt signaling[27, 28]. Silencing of AXIN2 could theoretically result in sustained activation of canonical Wnt/β-catenin signaling; however, its concurrent downregulation with WNT5A may reflect a more complex, tissue-specific rewiring of the Wnt network that favors bone resorption over formation. This apparent paradox warrants further dissection in future functional studies.

In contrast, the observed hypomethylation and increased expression of TGFB1 align with its complex, stage-dependent role in bone metabolism. Although TGFB1 is essential for early bone formation, its sustained overexpression in the bone microenvironment is a well-recognized driver of abnormal bone remodeling. It promotes the recruitment of osteoclast precursors and stimulates bone resorption, while ultimately leading to fibrous tissue deposition and impaired bone quality[29]. Thus, its epigenetic activation may be a key mechanism driving the high-turnover imbalance observed in some osteoporotic patients.

Similarly, the epigenetic upregulation of STAT3 links epigenetic alteration to inflammatory bone loss. Persistent STAT3 activation can promote osteoclast differentiation and survival while inhibiting osteoblast function, creating a vicious cycle that exacerbates net bone loss[30]. This positions STAT3 as a potential epigenetic mediator connecting chronic low-grade inflammation—often associated with aging—to the pathogenesis of osteoporosis.

Importantly, these genes do not function in isolation but act as nodes within interconnected pathways. The silencing of osteogenic transcription factors and specific Wnt components, coupled with the activation of catabolic signals, paints a coherent picture of a biased epigenetic program that disrupts bone homeostasis[31]. This network-level dysregulation suggests that targeting a single gene may be insufficient. Instead, the identified epigenetic marks themselves—particularly the reversible promoter hypermethylation of genes such as MSX1 and AXIN2—hold potential as novel therapeutic targets for demethylating agents or as diagnostic biomarkers for patient stratification[32].

While this integrative study provides novel insights into the epigenetic landscape of osteoporosis, several limitations must be acknowledged. First, the sample size of our discovery cohort, although utilizing precious bone biopsy specimens, remains modest, which may constrain the statistical power to detect more subtle methylation alterations. Second, the correlative nature of our bulk tissue analysis precludes the establishment of direct causality between the observed DNA methylation changes and downstream gene expression shifts or the osteoporotic phenotype. Third, the observational nature of our study, without direct experimental perturbation, limits causal inference. The functional impact of the prioritized epigenetic alterations on osteoblast or osteoclast activity warrants future validation using targeted in vitro or in vivo models. Finally, the clinical translatability of our findings necessitates validation in larger, independent, and well-characterized longitudinal cohorts to rigorously assess the prognostic value and generalizability of the identified epigenetic signatures.

Future research should extend these findings along several critical directions. For instance, applying single-cell multi-omics technologies to bone samples would enable the direct mapping of methylation and expression changes to specific cell types, thereby clarifying the precise cellular origins and impacts of the observed epigenetic dysregulation[33]. Subsequently, direct functional testing of our prioritized candidate genes, particularly MSX1 and AXIN2, is essential. Employing CRISPR-dCas9-based epigenetic editing tools to specifically reverse promoter hypermethylation at these loci in relevant bone cell models would be crucial for conclusively establishing their causal role in driving osteogenic dysfunction[34].

Nevertheless, our work delineates a coherent set of interconnected, epigenetically dysregulated genes and pathways. Further investigation along the proposed directions will not only solidify the pathophysiological understanding of epigenetic contributions to osteoporosis but will also facilitate the development of novel epigenetic diagnostics and targeted therapeutic strategies.

## Conclusion

This study employed whole-genome bisulfite sequencing to profile genome-wide DNA methylation differences between patients with osteoporosis and non-osteoporosis controls. The osteoporotic group exhibited a higher global methylation burden and a greater number of differentially methylated genes compared to the control group. Through integrated enrichment analysis and validation using public databases, we prioritized six genes—MSX1, HOXD4, AXIN2, WNT5A, TGFB1, and STAT3—as potential epigenetic drivers of osteoporosis, likely acting through distinct biological pathways. We acknowledge that the limited sample size and the multiple-gene screening approach may introduce bias in identifying robust differential methylation signatures; thus, these findings require further validation. Functional experiments are also necessary to test the mechanistic hypotheses generated from these methylome data. Nonetheless, this work contributes to the growing understanding of epigenetic dysregulation in osteoporosis and may inform the development of novel diagnostic and therapeutic strategies.

## Abbreviations

EWAS: Epigenome-wide association studies
DMRs: Differentially methylated regions
WGBS: Whole-genome bisulfite sequencing
BMD: Bone mineral density
BMI: Body mass index
CGI: CpG island
GO: Kyoto Encyclopedia of Genes and Genomes
KEGG: Kyoto Encyclopedia of Genes and Genomes
PPI: Protein-Protein Interaction
FDR: False discovery rate
DNMT: DNA Methyltransferase
RUNX2: Runt-related transcription factor 2
MSX1: Msh Homeobox 1
HOXD4: Homeobox D4
AXIN2: Axin 2
WNT5A: Wnt Family Member 5A
TGFB1: Transforming Growth Factor Beta 1
STAT3: Signal Transducer And Activator Of Transcription 3.

## Data availability

Sequence data that support the findings of this study have been deposited in the NCBI with the primary accession code PRJNA1154659

## Acknowledgements

We sincerely thank all members of Ying Li’s group for their contributions to sample collection and technical assistance. All images were created by the author team.

## Authors’ contributions

YYZ and GYW conducted the experiments and data analysis. JLH designed and supervised the experiment and assisted with manuscript revisions. YLZ provided technical support. YKZ and SSX coordinated the research. ZHG and YL performed the surgeries and provided the samples. YYZ drafted the manuscript. All authors read and approved the final version of the manuscript.

## Ethics approval

All experimental procedures, including human care and tissue sample collection, were approved by and conducted in accordance with the guidelines of the Medical Ethics Committee of the Third Affiliated Hospital of Guangzhou University of Chinese Medicine, China (Approval ID: PJ-XS-20240531-009).

## Consent for publication

Not applicable.

